# Combinatorial metabolic engineering of *Saccharomyces cerevisiae* for terminal alkene production

**DOI:** 10.1101/022350

**Authors:** Binbin Chen, Dong-Yup Lee, Matthew Wook Chang

## Abstract

Biological production of terminal alkenes has garnered a significant interest due to their industrial applications such as lubricants, detergents and fuels. Here, we engineered the yeast *Saccharomyces cerevisiae* to produce terminal alkenes via a one-step fatty acid decarboxylation pathway and improved the alkene production using combinatorial engineering strategies. In brief, we first characterized eight fatty acid decarboxylases to enable and enhance alkene production. We then increased the production titer 7-fold by improving the availability of the precursor fatty acids. We additionally increased the titer about 5-fold through genetic cofactor engineering and gene expression tuning in rich medium. Lastly, we further improved the titer 1.8-fold to 3.7 mg/L by optimizing the culturing conditions in bioreactors. This study represents the first report of terminal alkene biosynthesis in *S. cerevisiae*, and the abovementioned combinatorial engineering approaches collectively increased the titer 67.4-fold. We envision that these approaches could provide insights into devising engineering strategies to improve the production of fatty acid-derived biochemicals in *S. cerevisiae*.

## 1. Introduction

Global focus towards reducing petroleum footprint has led to a significant interest in developing alternative methods to produce fuels from low-cost and renewable resources. Metabolic engineering has emerged as an enabling technology to this end, which directs modulation of metabolic pathways by using recombinant technologies to overproduce valuable products, including biofuels (de Jong et al., 2011; Peralta-Yahya et al., 2012; Stephanopoulos, 2012; Zhang et al., 2011). Alkenes, traditionally used as detergents, lubricating fluids and sanitizers (Wang et al., 2013), have the potential to serve as “drop-in” compatible hydrocarbon fuels because of their high energy content. In addition, as they are already predominant components of petroleum-based fuels (Howard et al., 2013; Poirier and George, 1982), they are compatible with the existing engine platform and fuel distribution systems. Therefore, there is a strong economic and environmental demand for the development of bio-alkenes, which could be low-cost and environmentally sustainable, through metabolic engineering strategies.

The fatty acid biosynthesis pathway is ideally suited to provide biofuel precursors because of the high energy content in the precursors, and these fatty acid precursors can be converted into alkenes via naturally occurring metabolic pathways (Beller et al., 2010; Mendez-Perez et al., 2011; Rude et al., 2011; Sukovich et al., 2010a; Sukovich et al., 2010b). The first pathway involves a cytochrome P450 fatty acid decarboxylase - OleT_JE_ from *Jeotgalicoccus* sp. ATCC 8456 which directly decarboxylates free fatty acids to terminal alkenes (Rude et al., 2011). The second pathway employs a multi-domain polyketide synthase, found in the cyanobacterium *Synechococcus* sp. PCC 7002. This enzyme converts fatty acyl-ACP to terminal alkene via an elongation decarboxylation mechanism (Mendez-Perez et al., 2011). The third pathway produces long-chain internal alkenes (C24 - C31) by a head-to-head condensation of two acyl-CoA (or-ACP) thioesters followed by several reduction steps in *Micrococcus luteus* (Beller et al., 2010) and *Shewanella oneidensis* (Sukovich et al., 2010a; Sukovich et al., 2010b). Among these three pathways, the one-step fatty acid decarboxylation pathway is highly advantageous for alkene biosynthesis for the following two reasons. Firstly, the fatty acid synthesis pathway is feedback-inhibited by fatty acyl-CoA/ACP (Davis and Cronan, 2001; Ogiwara et al., 1978), a precursor of fatty acid-derived biofuels. This feedback inhibition could negatively affect the boosting of fatty acyl-CoA/ACP levels, and in turn the fatty acid-derived biofuel titers. Thus, using free fatty acids as biofuel precursors is more desirable compared with fatty acyl-CoA/ACP. Secondly, a one-step reaction from fatty acids to alkenes reduces intermediate metabolite losses and toxicity (Conrado et al., 2012; Kizer et al., 2008; Pitera et al., 2007).

The well-studied industrial microorganism *Saccharomyces cerevisiae* offers a number of advantages (Kalscheuer et al., 2004; Leber and Da Silva, 2013; Nevoigt, 2008; Yu et al., 2010) for producing fatty acid-derived products due to i) its ability to withstand lower temperatures, ii) immunity towards phage contaminations, iii) suitability in large-scale fermentation, iv) generally higher tolerance toward abiotic stresses, and v) extensive knowledge available about its fatty acid metabolism.

Consequently, in this study, we aimed to engineer the yeast *S. cerevisiae* to produce terminal alkenes via a one-step fatty acid decarboxylation pathway and to improve the alkene production using combinatorial engineering strategies (depicted in Figure 1A). First, we screened and characterized eight fatty acid decarboxylases (OleT) to enable and enhance alkene production in *S. cerevisiae*. We then developed a fatty acid-overproducing strain to boost the precursor availability, which could enhance the metabolic flux (Scalcinati et al., 2012) and resulted in a higher production titer. We then improved the enzyme cofactor accumulation through cofactor genetic engineering (Lopez de Felipe et al., 1998; Scalcinati et al., 2012). We then enhanced the cell growth in rich medium and tuned the enzyme expression by optimizing the combinations of the promoters and plasmids. Finally, we further increased the alkene production by optimizing the culturing conditions in bioreactors. This study represents the first report of terminal alkene biosynthesis in the yeast *S. cerevisiae*, and the abovementioned combinatorial engineering approaches collectively increased the titer of the alkene production of *S. cerevisiae* 67.4-fold.

**Fig. 1.**
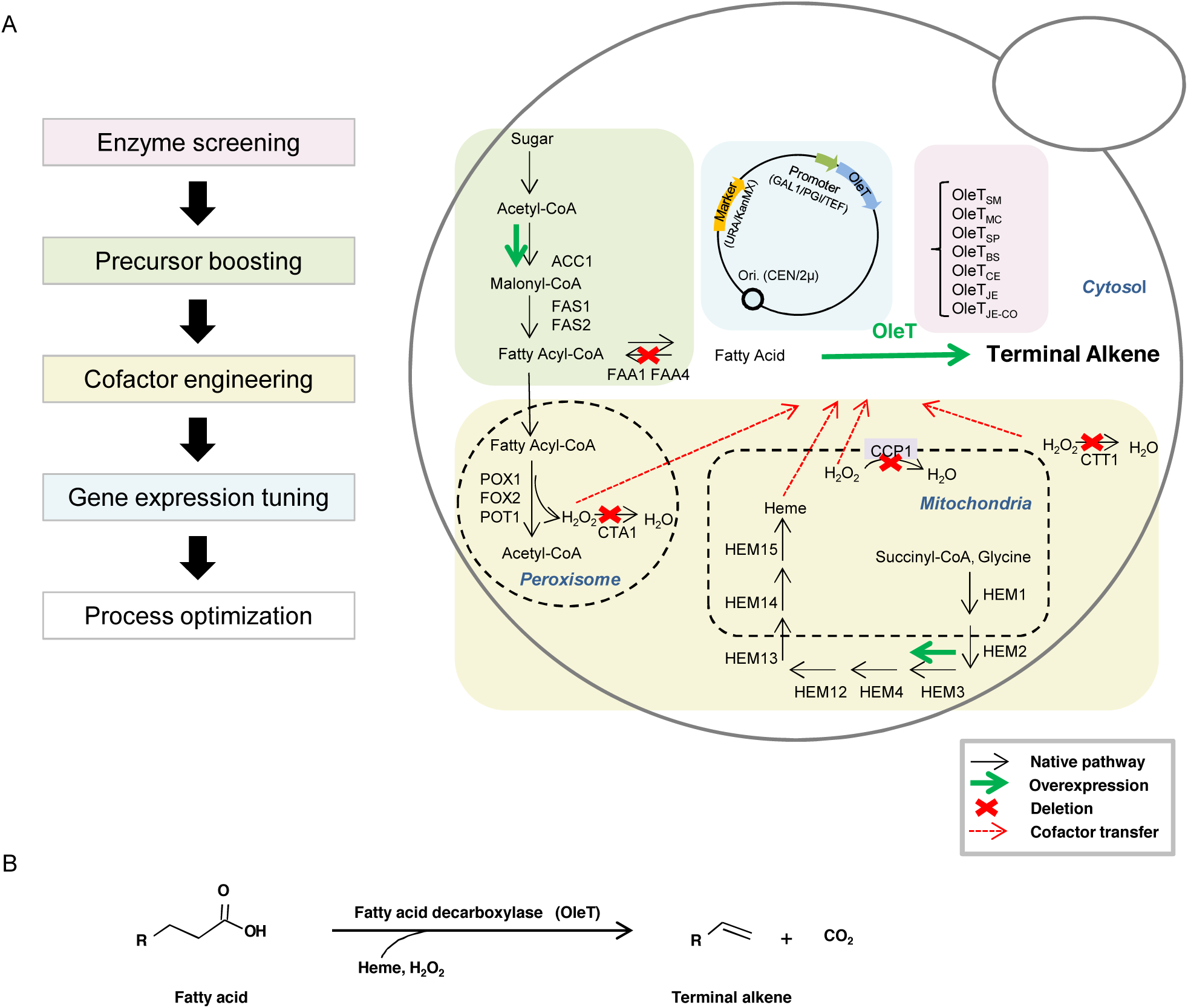
(A) Schematic view of the metabolic pathway for the production of terminal alkenes in the genetically engineered strain. Black arrows represent the native pathway in *S. cerevisiae*; Green arrows represent the overexpression of genes in this study; Red crosses represent the gene deletion performed. Red dash arrows represent cofactor transfer for OleT utilization. ACC1: acetyl-CoA carboxylase; FAS1/2: fatty acid synthase; FAA1/4: fatty acyl-CoA synthetase; POX1: fatty acyl-CoA oxidase; FOX2: 3-hydroxyacyl-CoA dehydrogenase and enoyl-CoA hydratase; POT1: 3-ketoacyl-CoA thiolase; CCP1: cytochrome c peroxidase; CTA1: catalase A; CTT1: catalase T; HEM1: 5-aminolevulinate synthase; HEM2: aminolevulinate dehydratase; HEM3: porphobilinogen deaminase; HEM4: uroporphyrinogen III synthase; HEM12: uroporphyrinogen decarboxylase; HEM13: coproporphyrinogen oxidase; HEM14: protoporphyrinogen oxidase; HEM15: ferrochelatase; OleT: fatty acid decarboxylase. (B) Synthesis of terminal alkene via fatty acid decarboxylase - OleT catalyzed reaction.

## 2. Materials and methods

### 2.1. Strains and media

*Escherichia coli* TOP10 (Invitrogen) and Luria-Bertani (BD) were used for cloning experiments unless otherwise stated. 100 mg/L ampicillin was used for selection of positive colonies if applicable. *Jeotgalicoccus* sp. ATCC 8456 (NCIMB) was used for *oleT*_*JE*_ cloning. The yeast strain *S. cerevisiae* BY4741 (ATCC) was used for functional characterization of OleT enzymes.

*S. cerevisiae* BY4741 wild-type and mutant strains were cultured in rich medium (YPD/YPG), synthetic minimal medium lacking uracil (SC-U), lysine (SC-L), adenine (SC-A), or synthetic minimal induction medium (SC-U-G). YPD/YPG medium (1% yeast extract, 2% peptone and 2% D-glucose/galactose) was used to routinely maintain wild-type strain or cells with pRS41K or pRS42K plasmids. SC-U medium (0.67% yeast nitrogen base, 0.192% uracil dropout and 2% raffinose) was used for growing pESC-URA transformants. SC-L medium (0.67% yeast nitrogen base, 0.18% lysine dropout and 2% glucose) and SC-A medium (0.67% yeast nitrogen base, 0.078% adenine dropout and 2% glucose) was used for selecting positive integrants. SC-U-G medium (0.67% yeast nitrogen base, 0.192% uracil dropout, 1% raffinose and 2% galactose) was used for protein induction in pESC-URA transformants. 2% agar was supplemented for solid media. 1mg/mL 5-Fluoroorotic acid (5-FOA, Fermentas) or 200 mg/L geneticin (G418, PAA Laboratories) was used for selection. Heme (20ug/mL) (Cho and Jeffries, 1999; Villarreal et al., 2008), hydrogen peroxide (0.4mM every 12 h) (Izawa et al., 1996), or both were supplemented into growth culture where necessary. Yeast growth media components were purchased from Sigma-Aldrich and MP Biomedicals. Yeast cells were cultivated at 30 °C in flasks and shaken at 250 rpm.

### 2.2 Gene deletion and integration

Genes were deleted by using the previously described gene disruption cassette containing loxP-kanMX-loxP module in *S. cerevisiae* (Güldener et al., 1996). Firstly, the gene disruption cassettes were constructed through fusing short homologous sequences with loxP-kanMX-loxP module from plasmid pUG6 (Euroscarf) via a PCR reaction. Following yeast transformation, colonies were selected on an YPD plate containing 200 mg/L G418. The kanMX marker was removed by inducing expression of Cre recombinase from plasmid pSH47 (Euroscarf), which enables subsequent rounds of gene deletion. Here, the correct gene deletion mutants were verified by PCR analysis and used for further gene deletion.

Chromosomal integration was conducted based on the method previously reported by Sadowski et al. (Sadowski et al., 2007). Briefly, genes were firstly cloned into plasmid pIS385 or pIS112 (Euroscarf) containing URA3 selectable marker. The recombinant plasmid was linearized and transformed into *S. cerevisiae*, followed by colony selection performed on SC-U medium. After non-selective growth on YPD plate, individual colonies were replica-plated onto 5-FOA and SC-L or SC-A plates to screen for positive colonies. Finally, the correct integrant was verified by PCR analysis. Oligonucleotide primers used for gene deletion and chromosomal integration are listed in Table S1 (Supplementary material).

### 2.3 Fatty acid decarboxylase selection

Six more homologous enzymes from different organisms were selected for alkene biosynthesis in *S. cerevisiae* (Table S2, Supplementary material). Among them, *oleT*_*BS*_, *oleT*_*MP*_ and *oleT*_*CE*_ were reported to produce 1-pentadecene when heterologously expressed in *E. coli* (Rude et al., 2011); *oleT*_*SM*_, *oleT*_*MC*_ and *oleT*_*SP*_ were selected based on their protein sequence identity to *oleT*_*JE*_, and their histidine residue in position 85 (His85) which as mentioned, plays an important role in catalysis activity of OleT_JE_.

### 2.4 Plasmid construction

To clone *oleT*_*JE*_, genomic DNA of *Jeotgalicoccus* sp. ATCC 8456 was used as a PCR template performed with two primers OleT_JE_-F and OleT_JE_-R. One *oleT*_*JE*_ codon optimized gene and six codon optimized *oleT*_*JE*_ homologous genes, namely *oleT*_*JE-CO*_, *oleT*_*SM*_, *oleT*_*MC*_, *oleT*_*SP*_, *oleT*_*BS*_, *oleT*_*MP*_, and *oleT*_*CE*_, were synthesized from Life technologies. *ACC1* and *HEM3* were amplified from *S. cerevisiae* genome using two set of primers: ACC1-SC-F and ACC1-SC-R, Hem3-F and Hem3-R. A list of primers used was shown in Table S1 (Supplementary material). Plasmid pESC-URA (Agilent Technologies), pRS41K (Euroscarf) and pRS42K (Euroscarf) were used as expression vectors for *oleT* and/or *ACC1* while plasmid pIS385 (Euroscarf) was used for *HEM3* cloning. Either Gibson DNA assembly method (Gibson et al., 2009) or digestion-ligation method was used for the construction of all the plasmids. The constructed recombinant plasmids are listed in Table 1.

**Table 1.**
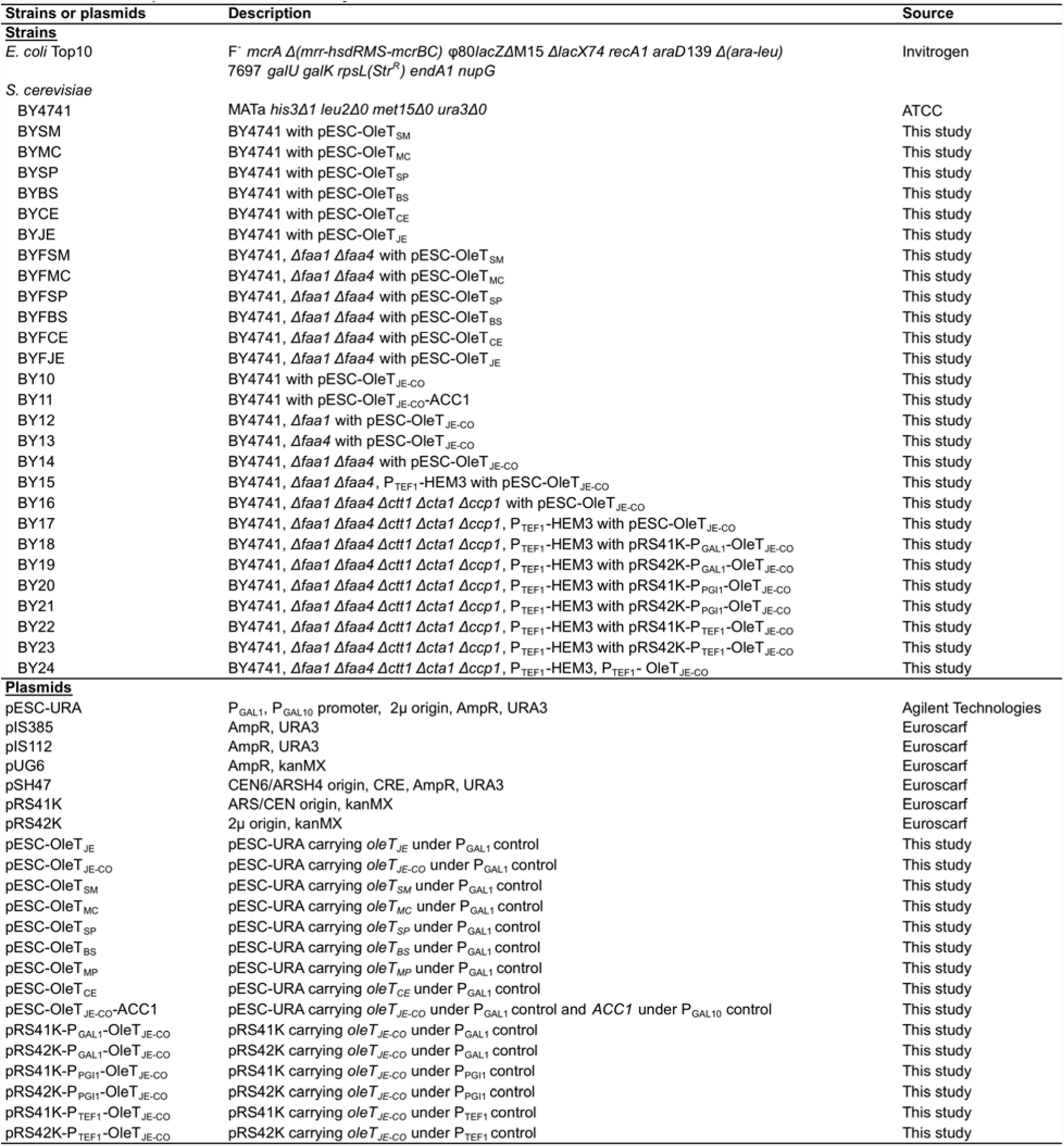
Strains and plasmids used in this study

### 2.5 Alkene extraction and detection

For alkene production, cells were pre-cultured in 10 ml medium overnight and then diluted in 50 ml induction medium using 250 ml flask to achieve an initial OD_600_ of 0.4. After growing for 48 h, yeast cells were harvested by centrifugation at 6000 g for 5 min. Cell pellets were re-suspended in HPLC grade methanol (Sigma), and 1-nonene was added into cell suspension as an internal standard. Acid-washed glass beads were added until the suspension was covered. Cells were then lysed by mechanical agitation using FastPrep-24 (MPBio) for 8 min at 6 m/s. HPLC grade hexane (Sigma) was then added and mixed thoroughly with crude extract for 5 min. The crude extract was separated into two phases by centrifugation, and the upper phase containing alkene was transferred into a clear GC vial.

The alkenes dissolved in the upper layer were quantified using gas chromatography-mass spectrometry (GC-MS) under the following conditions. An HP-5ms column (30 m by 0.25 mm; 0.25 μm film; Agilent) was used with a helium flow rate set to 1.1 ml/min. Injections of 5 μl were carried out under splitless injection condition with the inlet set to 250 °C. The GC temperature profile was as follows: an initial temperature of 40 °C was maintained for 0.5 min, followed by ramping to 280 °C at a rate of 6 °C/min, where the temperature was held for 3 min. The mass spectrometer detector was scanned at 30 to 800 amu in the electron impact mode. To aid peak identification, authentic references (C9-C19 terminal alkenes, Tokyo Chemical Industry) were used, and their retention times and fragmentation patterns were compared with those from the extracted alkenes.

### 2.6 Bioreactor conditions

Selected strain was used for production of alkenes through fed-batch fermentation. YPD+G418 containing 3% glucose was used for both seed preparation and fermentation. Seed culture was prepared by inoculating colonies into a 250 mL flask containing 50 mL culture medium, and incubating at 30 °C and 250 rpm for 24 h. The seed was then transferred to a 5 L bioreactor (BIOSTAT® B-DCU II, Sartorius) containing 1 L medium with an initial OD_600_ 0.4. The fermentation was carried out at 30 °C. The dissolved oxygen concentration in the bioreactor was maintained at around 60% by controlling the air flow rate and agitation speed. 150ml 200 g/L glucose was fed to the fermenter every 24h and samples were withdrawn at the indicated time intervals. All of the fermentation experiments were performed in triplicate.

## 3. Results and discussion

### 3.1. Screening enzymes for alkene biosynthesis in *S. cerevisiae*

To enable terminal alkene production in *S. cerevisiae*, we attempted to use the cytochrome P450 fatty acid decarboxylase OleT_JE_ from *Jeotgalicoccus* sp. ATCC 8456, which reportedly decarboxylates fatty acids to terminal alkenes (Rude et al., 2011) (Figure 1B). We also used its codon-optimized version *oleT*_*JE-CO*_ and six of its homologous genes, based on high sequence identity to OleT_JE_ (Table S2, Supplementary material). Native *oleT*_*JE*_ and synthesized codon-optimized homologous genes were cloned into the high copy plasmid pESC-URA (Table 1) and transformed into *S. cerevisiae*. The induced protein expression in *S. cerevisiae* was confirmed by western blot (data not shown). We evaluated the performance of the abovementioned enzymes by quantifying the alkene profiles and measuring the alkene concentrations from the cell cultures grown for 48 h. We found that the cells carrying the empty plasmid and OleT_MP_ from *Methylobacterium populi* BJ001 produced no detectable alkenes (data not shown), whilst the transformants expressing the other OleT enzymes produced a range of alkenes. As shown in Figure 2, OleT_SM_, OleT_SP_, OleT_BS_ and OleT_CE_ produced alkenes with the chain lengths of C13, C15 and C17, whereas OleT_MC_ exhibited a narrower alkene profile, producing C13 and C15 alkenes. OleT_JE_ and its codon-optimized version OleT_JE-CO_ exhibited the broadest product profile range, producing odd chain terminal alkenes from C11 to C19. We observed lower alkene titers for shorter chain lengths possibly because longer chain fatty acids are more abundant than shorter chain fatty acids in yeast cells (Welch and Burlingame, 1973).

**Fig. 2.**
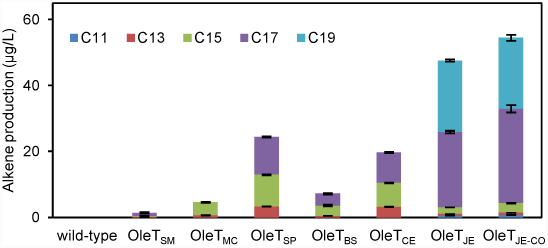
Production of alkenes by recombinant *S. cerevisiae* expressing *oleT*_*JE*_ homologs. Distributions of different chain length alkenes produced by the overexpression of *oleT*_*SM*_, *oleT*_*MC*_, *oleT*_*SP*_, *oleT*_*BS*_, *oleT*_*CE*_, *oleT*_*JE*_ and *oleT*_*JE-CO*_ are shown. Alkenes with different chain lengths from C11 to C19 are represented with different colors. Results are the average of three biological replicates with error bars showing the standard deviation from the mean value.

Aside from the varying alkene profiles, the total titers of the produced alkenes varied among the tested OleT enzymes. Figure 2 shows that OleT_SM_ led to the lowest total alkene titer (1.4 μg/L), whereas OleT_JE-CO_ gave the highest total alkene titer (54.5 μg/L), which served as the baseline titer for this study.

### 3.2. Increase in free fatty acid production improved alkene production

As a first step in improving the alkene production, we attempted to increase the production of free fatty acids, which are precursors to alkenes (Figure 1A). The *de novo* fatty acid biosynthesis in *S. cerevisiae* requires acetyl-CoA carboxylase (ACC1; encoded by the *ACC1*) and fatty acid synthase complex (FAS; encoded by *FAS1* and *FAS2*) (Ruenwai et al., 2009; Runguphan and Keasling; Shin et al., 2012; Tai and Stephanopoulos, 2013). ACC1 converts acetyl-CoA into malonyl-CoA, and the overexpression of *ACC1* results in increase in final fatty acid level (Ruenwai et al., 2009; Tai and Stephanopoulos, 2013). The FAS complex produces fatty acyl-CoAs by condensation of one acetyl-CoA to 7∼8 malonyl-CoAs (Tehlivets et al., 2007). The *de novo* produced fatty acyl-CoAs are further hydrolyzed to free fatty acids; however, free fatty acids are converted back to fatty acyl-CoAs by endogenous fatty acyl-CoA synthetase (FAA1-4, FAT1). As an active form of free fatty acids, fatty acyl-CoAs are further degraded mainly through β-oxidation pathway (POX1, FOX2, POT1). Hence, in order to enhance the metabolic flux towards free fatty acid, we attempted to overexpress *ACC1* and disrupt *FAA1* and *FAA4*, the two main fatty acyl-CoA synthetases (Black and DiRusso, 2007; Choi and Martin, 1999; Scharnewski et al., 2008).

First, we expressed *oleT*_*JE-CO*_ with *ACC1* under the control of the strong inducible promoters P_GAL1_ and P_GAL10_, respectively, generating the strain BY11 (*ACC1, oleT*_*JE-CO*_). Second, we deleted *FAA1* and/or *FAA4*, and expressed *oleT*_*JE-CO*_, resulting in three different strains BY12 (*Δfaa1, oleT*_*JE-CO*_), BY13 (*Δfaa4, oleT*_*JE-CO*_), and BY14 (*Δfaa1Δfaa4, oleT*_*JE-CO*_). As shown in Figure 3A, the co-expression of *ACC1* and *oleT*_*JE-CO*_ in *S. cerevisiae* led to lower alkene levels compared with the singularly expressed *oleT*_*JE-CO*_ (BY10, control strain). Moreover, increased alkene production levels were observed in both BY12 (6.2-fold) and BY14 (7-fold). In particular, the double-deletion strain BY14 produced the highest alkene titer of 382.8 μg/L. However, for an unknown reason, the single-deletion of *FAA4* (BY13) led to 2.5-fold lower alkene production. These results suggest that the deletion of *FAA1* in tandem with *FAA4* has a synergic effect on fatty acid accumulation, where *FAA1* accounts for most of this effect. In addition to the total alkene titers, changes in alkene profiles were also studied (Table 2, Figure 3B). As shown in the gas chromatography (GC) profile, BY14 showed a significant improvement in the production of C15 and C17 alkenes compared to BY10, but a lower improvement for other alkenes. This increase in the production of C15 and C17 alkenes could be attributed to that BY14 accumulated more C16 and C18 free fatty acids (data not shown). We then expressed all eight OleT enzymes in the double-deletion strain (*Δfaa1Δfaa4*), respectively, and evaluated the alkene titers. We found that the overexpression of *oleT*_*JE-CO*_ showed the highest total alkene titer in the double-deletion strain (*Δfaa1Δfaa4*) (Figure 3C), in line with the result from the overexpression of *oleT*_*JE-CO*_ in the wild-type strain. Thus, we selected BY14 (*Δfaa1Δfaa4, oleT*_*JE-CO*_) for further engineering, which showed a 7-fold improvement in the titer to the control alkene-producing strain BY10 (*oleT*_*JE-CO*_).

**Fig. 3.**
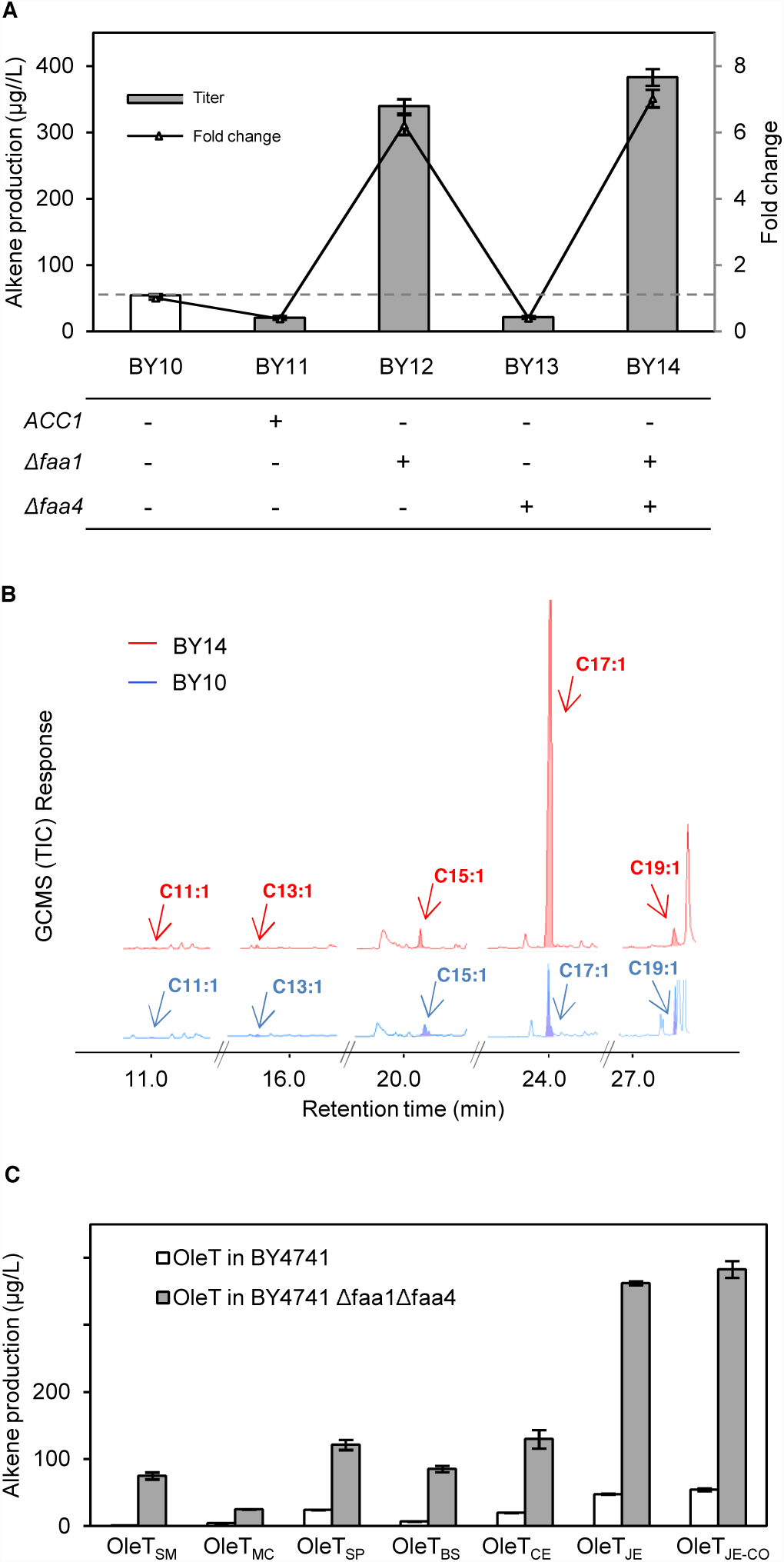
Effects of fatty acid pool engineering on alkene production. (A) Total alkene titers of the strains without (BY10) and with the engineered fatty acid synthesis pathway (BY11, BY12, BY13 and BY14) are shown in bars. White bar and grey horizontal dash line indicates the alkene titers of the control strain BY10. Alkene fold changes are shown in lines. For alkene fold changes, BY10 was set equal to 1.0 and all values were determined relative to BY10. “+” and “-” indicate with and without engineering respectively. (B) Gas chromatography (GC) profile of the alkene products obtained by batch culture of BY14 (red line) and BY10 (blue line). Filled peaks indicated by arrows were shown as specific alkenes. (C) The comparison of total alkenes produced by the expression of *oleT*_*JE*_ homologs in wild-type BY4741 (white bar) and BY4741 *Δfaa1Δfaa4* double-deletion strain (grey bar). Alkenes were detected and quantified by GC-MS after growing for 48 h. Results represent the mean of three biological replicates; standard deviations are presented.

**Table 2:** Comparison of alkene production obtained by engineered *S. Cervisiae* strains.

| Strain | Alkene (fractional abundance %) | | | | | Total alkene<br>($\mu\text{g/L}$) |
| --- | --- | --- | --- | --- | --- | --- |
|  | C11 | C13 | C15 | C17 | C19 |  |
| BYSM | - | 17.3 | 10.3 | 72.4 | - | 1.4±0.3 |
| BYMC | - | 14.0 | 86.0 | - | - | 4.6±0.1 |
| BYSP | - | 13.7 | 39.3 | 46.9 | - | 24.4±0.3 |
| BYBS | - | 6.5 | 44.2 | 49.3 | - | 7.2±0.4 |
| BYCE | - | 16.3 | 36.8 | 46.8 | - | 19.7±0.3 |
| BYJE | 1.2 | 1.0 | 4.2 | 48.0 | 45.6 | 47.6±0.8 |
| BYFSM | - | 2.6 | 13.1 | 84.3 | - | 75.0±5.2 |
| BYFMC | - | 7.3 | 92.7 | - | - | 25.5±0.3 |
| BYFSP | - | 3.1 | 41.1 | 55.8 | - | 121.2±7.6 |
| BYFBS | - | 3.3 | 33.4 | 63.3 | - | 85.4±4.7 |
| BYFCE | - | 2.3 | 36.8 | 61.0 | - | 129.6±13.7 |
| BYFJE | 0.2 | 0.5 | 5.0 | 88.2 | 6.1 | 362.1±3.0 |
| BY10 | 1.4 | 1.3 | 5.4 | 52.4 | 39.5 | 54.5±2.2 |
| BY11 | - | 2.6 | - | 44.4 | 52.9 | 21.0±2.3 |
| BY12 | - | 0.3 | 2.2 | 94.1 | 3.5 | 339.2±10.8 |
| BY13 | - | 2.7 | 4.9 | 46.5 | 45.8 | 21.8±2.0 |
| BY14 | 0.2 | 0.4 | 4.1 | 89.5 | 5.9 | 382.8±12.6 |
| BY14<sup>a</sup> | 0.3 | 0.8 | 7.0 | 87.6 | 4.2 | 716.9±30.0 |
| BY14<sup>b</sup> | 1.0 | 1.6 | 14.0 | 79.1 | 4.3 | 684.0±27.5 |
| BY14<sup>c</sup> | 0.4 | 1.2 | 8.5 | 87.0 | 3.0 | 1387.4±48.9 |
| BY15 | 0.1 | 0.3 | 3.7 | 92.2 | 3.7 | 403.8±5.4 |
| BY16 | 0.1 | 0.4 | 4.3 | 91.1 | 4.2 | 402.0±13.9 |
| BY17 | 0.2 | 0.3 | 3.1 | 92.5 | 3.8 | 472.7±8.6 |
| BY18 | 0.2 | 1.1 | 8.3 | 77.2 | 13.2 | 1720.8±156.9 |
| BY19 | 1.0 | 1.4 | 9.0 | 69.6 | 19.0 | 453.2±29.8 |
| BY20 | 0.6 | 1.0 | 10.7 | 71.9 | 15.8 | 409.9±25.9 |
| BY21 | 0.3 | 0.5 | 10.2 | 77.9 | 11.1 | 882.2±195.4 |
| BY22 | 0.1 | 0.4 | 8.3 | 85.0 | 6.2 | 2088.7±66.4 |
| BY23 | 0.6 | 0.9 | 9.5 | 59.6 | 29.4 | 551.9±16.3 |
| BY24 | 0.6 | 0.8 | 8.7 | 70.2 | 19.7 | 450.8±3.8 |
| BY22<sup>d</sup> | 0.4 | 0.3 | 5.8 | 52.1 | 41.4 | 763.9±32.4 |
| BY22<sup>e</sup> | 1.0 | 0.3 | 4.4 | 82.0 | 12.2 | 2243.5±117.3 |
| BY22<sup>f</sup> | 0.2 | 0.6 | 3.9 | 74.5 | 20.8 | 3289.1±217.9 |
| BY22<sup>g</sup> | 0.6 | 0.7 | 5.6 | 58.6 | 34.5 | 3675.5±218.4 |

a: Heme supplementation in medium
b: H<sub>2</sub>O<sub>2</sub> supplementation in mediumc: Heme and H<sub>2</sub>O<sub>2</sub> supplementation in medium
d: 24h growth in bioreactor
e: 48h growth in bioreactor
f: 72h growth in bioreactor
g: 144h growth in bioreactor
- not detected

### 3.3 Cofactor engineering further increased alkene production level

#### 3.3.1 Supplementation of cofactors: heme and hydrogen peroxide

We then improved the enzyme cofactor availability to further increase the associated metabolic flux towards alkene production. OleT_JE_ is a cytochrome P450 enzyme in the cyp152 family, which contains heme as a cofactor (Rude et al., 2011), and the overexpression of cytochrome P450 enzymes can lead to heme depletion (Michener et al., 2012). Further, OleT_JE_ is highly active in the presence of hydrogen peroxide which serves as the sole electron and oxygen donor (Rude et al., 2011). Therefore, we hypothesized that cellular depletion of heme and hydrogen peroxide resulting from the overexpression of the P450 enzyme OleT_JE_ could be a limiting factor, and thus, increasing the availability of the two cofactors heme and hydrogen peroxide might improve alkene synthesis.

To test this hypothesis, we supplemented BY14 (*Δfaa1Δfaa4, oleT*_*JE-CO*_) with heme, hydrogen peroxide, or both. As shown in Figure 4, the supplementation with heme, hydrogen peroxide or both increased the titer by 87%, 79%, and 3.6-fold respectively, with the highest production at 1.4 mg/L. The improved alkene production demonstrated that cofactors supplementation during OleT enzyme expression could be employed to boost the alkene titers.

**Fig. 4.**
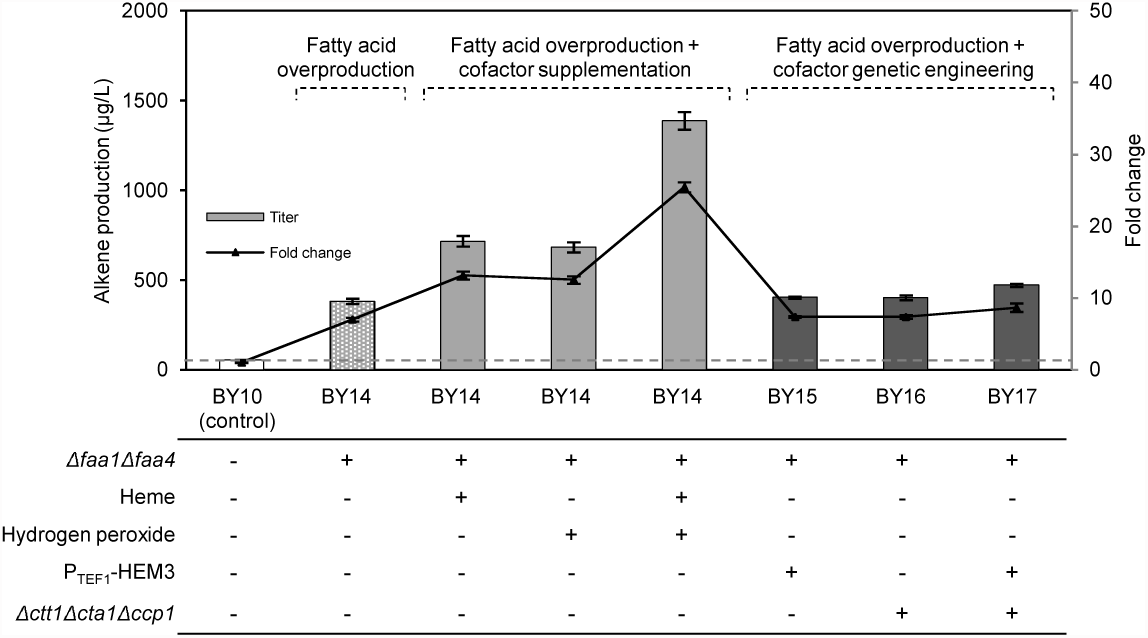
Production of alkenes by cofactor engineering. Total alkene titers are shown in bars and alkene fold changes are shown in lines. White bars and grey horizontal dash lines indicate the alkene titers of the control strain BY10. Lattice bars represent samples with fatty acid overproduction; Grey color bars represent sampless with fatty acid overproduction and cofactor supplementation; Black color bars represent samples with fatty acid overproduction and cofactor genetic engineering. For alkene fold changes, BY10 was set equal to 1.0 and all values were determined relative to BY10. “+” and “-” indicate with and without engineering respectively. Error bars represent the standard deviation of three biological replicates

#### 3.3.2 Overexpression of *HEM3*, and triple-deletion of *CTT1*, *CTA1* and *CCP1*

Based on the abovementioned result from the cofactor supplementation, we attempted to increase the alkene titer using genetic cofactor engineering to eliminate the need for cofactor supplementation, which could be costly. We first aimed to improve cellular heme production, which could be achieved by overexpression of rate-limiting enzymes responsible for heme biosynthesis. Multiple enzymes are involved in the heme biosynthesis pathway including three rate-limiting enzymes, HEM2, HEM3 and HEM12 (Hoffman et al., 2003); however, the co-expression of these three HEM enzymes could be stressful to the host cells (Michener et al., 2012). For example, the strains expressing only *HEM3* exhibited no growth defect, and in combination with expression of P450 enzyme, showed high theophylline titers (Michener et al., 2012). Therefore, in this study, *HEM3* was integrated into genome and constitutively expressed under the control of *TEF1* promoter, referred to as strain BY15 (*Δfaa1Δfaa4, P*_*TEF1*_*-HEM3, oleT*_*JE-*_ *CO*). Secondly, we aimed to accumulate endogenous hydrogen peroxide by deleting its utilization enzymes, catalase T (CTT1) located in cytoplasm, catalase A (CTA1) located in peroxisomes (Petrova et al., 2002), and the antioxidant enzyme cytochrome c peroxidase (CCP1) located in mitochondria (Verduyn et al., 1988). Previous studies showed that increased levels of hydrogen peroxide were detected in catalase mutants and cells with chemically inactivated catalases (Mesquita et al., 2010; Zhang and Wang, 1994). Hence, we further deleted *CTT1*, *CTA1* and *CCP1* genes to generate a series of deletion strains that could improve cofactor availability (Table 1).

As shown in Figure 4, *HEM3* expression (BY15) brought a slight improvement in the total alkene titer compared to BY14 (without *HEM3* overexpression). However, among all the deletion mutants, only BY16 (*Δfaa1Δfaa4Δctt1Δcta1Δccp1, oleT*_*JE-CO*_) showed a slightly higher titer compared to BY14 (*Δfaa1Δfaa4, oleT*_*JE-CO*_), while the rest deletion mutants showed no improved alkene titers (data not shown). To examine the potential synergistic effect of the aforementioned two approaches, we integrated *HEM3* into the genome of BY16, resulting in BY17 (*Δfaa1Δfaa4Δctt1Δcta1Δccp1, P*_*TEF1*_*-HEM3, oleT*_*JE-*_ *CO*). As shown in Figure 4, BY17 (*Δfaa1Δfaa4Δctt1Δcta1Δccp1, P*_*TEF1*_*-HEM3, oleT*_*JE-CO*_) produced a total alkene tilter of 472.7 μg/L, 23% improvement to the fatty acid-overproducing strain BY14 (*Δfaa1Δfaa4, oleT*_*JE-CO*_) and 8.7-fold improvement to the control strain BY10 (*oleT*_*JE-CO*_).

#### 3.4 Gene expression tuning for alkene production in rich medium

We then enhanced the cell growth in rich medium and tuned the expression level of the heterologous genes. In the highest producing strain so far BY17, the *oleT*_*JE-CO*_ was placed under the control of the galactose inducible promoter P_GAL1_ on the high-copy plasmid pESC-URA containing the auxotrophic URA marker. Rich medium frequently increase cell growth and final cell amount, resulting in higher product titers (Hahn-Hagerdal et al., 2005). Thus, here we replaced the auxotrophic pESC-URA plasmid with pRS plasmids containing the KanMX resistance marker. Moreover, to optimize the expression level of the heterologous genes, we used pRS41K (low copy) and pRS42K (high copy) as cloning vectors (Wang et al., 2012). P_GAL1_ (a strong inducible promoter), P_PGI1_ (a weak constitutive promoter) and P_TEF1_ (a strong constitutive promoter) were employed in both vectors to modulate the *oleT*_*JE-CO*_ transcription. A total of six engineered strains were constructed and tested for alkene production (Table 1).

All the engineered *oleT*_*JE-CO*_ containing strains were cultivated in rich medium supplied with 2% galactose or glucose for alkene production. We found that all the engineered strains exhibited increased cell growth and much higher final cell amount, where OD_600_∼30 was achieved in the rich medium while OD_600_∼8 in the minimal medium). As shown in Figure 5, among the abovementioned six constructed strains, BY22 (*Δfaa1Δfaa4Δctt1Δcta1Δccp1, P*_*TEF1*_*-HEM3, P*_*TEF1*_*-oleT*_*JE-CO*_ *(pRS41K)*), which contains the strong constitutive promoter P_TEF1_ on the low copy plasmid pRS41K, showed the highest alkene production, 2.1 mg/L, 4.4-fold higher than BY17 and 38.3-fold higher than the control strain BY10. The strains containing *oleT*_*JE-CO*_ under the control of the weak promoter P_PGI1_ showed 2.2-fold higher alkene production on the high copy plasmid pRS42K (BY21) than that on the low copy plasmid pRS41K (BY20). This result indicates that sufficient expression of *oleT*_*JE-CO*_ is needed for relatively higher alkene production. In contrast, with the strong promoter P_GAL1_ or P_TEF1_, the strains with the high copy plasmid (BY19 and BY23) showed 3.8-fold lower alkene production compared with the strains with the low copy plasmid (BY18 and BY22). These results suggest that in our study, i) the use of a strong promoter on a low copy plasmid provided sufficient enzyme levels for alkene production and ii) the use of a strong promoter on a high copy plasmid might cause “metabolic burden” on the cell, making the overall process non-beneficial (Ostergaard et al., 2000). To further address the “plasmid burden” (Karim et al., 2013) and to avoid the antibiotics cost, *oleT*_*JE-CO*_ was chromosomally integrated and constitutively expressed under the *TEF1* promoter. This constructed strain BY24 (*Δfaa1Δfaa4Δctt1Δcta1Δccp1, P*_*TEF1*_*-HEM3, P*_*TEF1*_*-oleT*_*JE-CO*_) produced about 4.6-fold less alkene than BY22 harboring *oleT*_*JE-CO*_ on a low-copy plasmid, suggesting that a single copy of *oleT*_*JE-CO*_ likely brought about insufficient gene expression level.

**Fig. 5.**
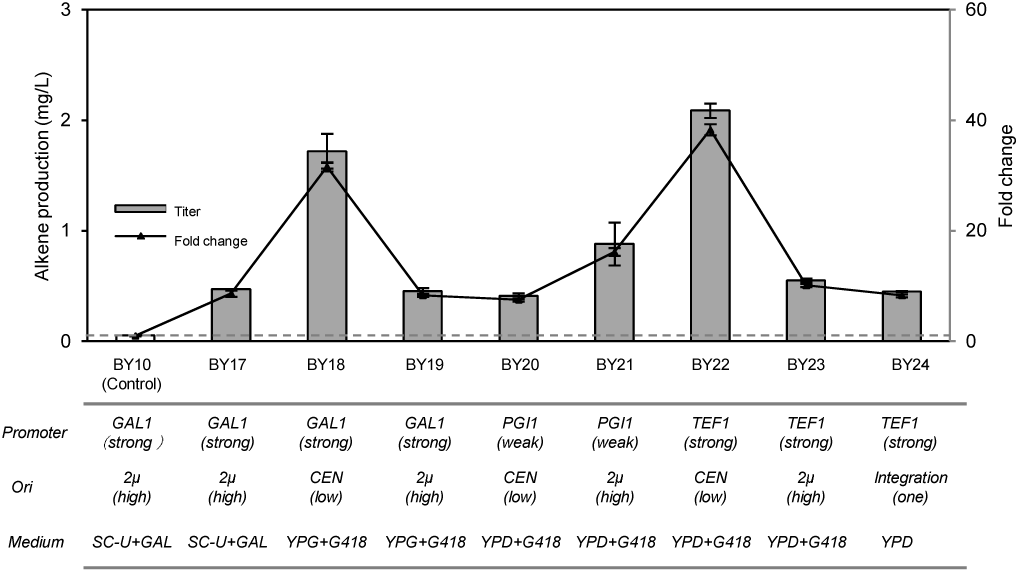
Alkene production using strains with tuned gene expression in rich medium. Total alkene titers are shown in bars and alkene fold changes are shown in lines. White bar and grey horizontal dash line indicates the alkene titers of control strain BY10. For alkene fold change, BY10 was set equal to 1.0 and all values were determined relative to BY10. Promoter strengths, plasmid copy numbers and respective growth medium are listed for each sample. Data shown are the mean ± SD of three biological replicates.

#### 3.5 Bioreactor process optimization for higher alkene production

We then conducted fed-batch fermentation and optimized the fermentation conditions to achieve higher alkene production. We used BY22 (*Δfaa1Δfaa4Δctt1Δcta1Δccp1, P*_*TEF1*_*-HEM3, P*_*TEF1*_*-oleT*_*JE-CO*_ *(pRS41K)*), the highest alkene production strain so far in shake flask culture, to test in fed-batch bioreactors. Three parameters, temperature, pH and dissolved oxygen concentration (pO2), were controlled and monitored. Three different operation temperatures, 25 °C, 30 °C and 35 °C gave comparable alkene titers (data not shown). pH 5, pH 7 and pH off were tested, where pH off showed a higher alkene titer (data not shown). Since heme biosynthesis requires oxygen (Hoffman et al., 2003) and an aerobic condition could give higher cell growth, the pO2 level was maintained at around 60% saturation, a general aerobic condition for yeast growth. Thus, we chose temperature 30 °C, pH off and pO_2_ 60% as our operation condition.

As shown in Figure 6A, during the first 48 h, BY22 grew steadily and the levels of the produced alkene were increased to 2.2 mg/L. After 48 h, strain went through the stationary phase, and the alkene levels were further increased from 2.2 mg/L to 3.3 mg/L at 72h; however, longer incubations only marginally increased alkene levels. These growth conditions gave rise to the highest level of production at 144 h, resulting in the alkene titer of 3.7 mg/L, 1.8-fold increase to the shake flask condition and 67.4-fold increase to the control strain BY10. Finally, Figure 6B and Table 2 summarize the abovementioned sequential improvements in the alkene production through enzyme screening, precursor boosting, cofactor engineering, gene expression tuning and process optimization.

**Fig. 6.**
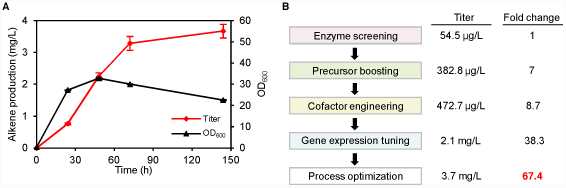
(A) Production of alkenes and cell optical density in 1-L fed-batch fermentation using the engineered strain BY22. Samples were withdrawn and analyzed at the indicated time intervals. Red lines indicate alkene titers and black line indicate cell OD. All of the fermentation experiments were performed in triplicate. (B) Titer and fold change summary for alkene production in *S. cerevisiae*.

## 4. Conclusions

In this study, we engineered *S. cerevisiae* to produce terminal alkene and further improved the alkene production 67.4-fold by combinatorial engineering strategies. First, OleT_JE_ and its homologous enzymes were characterized to convert free fatty acids into alkenes. In particular, OleT_JE-CO_ (codon optimized OleT from *Jeotgalicoccus* sp.) showed the broadest alkene profiles and the highest production level. Second, the deletion of both *FAA1* and *FAA4* significantly improved the alkene titer, likely due to increased free fatty acid pool. Third, genetic cofactor engineering involving the overexpression of *HEM3* and the triple-deletion of *CTT1*, *CTA1* and *CCP1* further improved the alkene titer. Fourth, the tuning of the heterologous gene expression in the rich medium enabled a further improvement in the titer (i.e. BY22 (*Δfaa1Δfaa4Δctt1Δcta1Δccp1, P*_*TEF1*_*-HEM3, P*_*TEF1*_*-oleT*_*JE-CO*_ *(pRS41K)*). Finally, the optimization of the culturing conditions in fed-batch bioreactors further improved the alkene production in BY22. This study represents the first report of terminal alkene biosynthesis in the yeast *S. cerevisiae*, and taken together, the abovementioned combinatorial engineering approaches increased the titer of the alkene production of *S. cerevisiae* 67.4-fold. We envision that these approaches could provide insights into devising engineering strategies to improve the production of fatty acid-derived biochemicals in *S. cerevisiae*.

## Acknowledgements

This work was supported by the Competitive Research Programme of the National Research Foundation of Singapore (NRF-CRP5-2009-03).

